# Excitation–inhibition interactions mediate firefly flash synchronization

**DOI:** 10.64898/2026.01.19.700439

**Authors:** Owen Martin, Nataliya Nechyporenko, Kaushik Jayaram, Orit Peleg

## Abstract

Large populations of fireflies can synchronize their bioluminescent flashes with remarkable precision, producing collective rhythms that emerge from interactions among intrinsically variable individuals. In the North American firefly *Photuris frontalis*, this behavior usually manifests as a stable, population-level single-period beat whose mechanistic origins remain unresolved. To identify the local interaction rules giving rise to this emergent synchrony, we performed controlled perturbation experiments on isolated *P. frontalis* males using fixed-period light stimuli. By measuring changes in flash period as a function of stimulus timing, we reconstructed the phase-response curve (PRC) governing individual flash dynamics. The resulting PRC exhibits a biphasic structure, revealing phase-advancing (excitatory) and phase-delaying (inhibitory) responses. Using this PRC, we formalized an integrate-and-fire model that quantitatively reproduced the observed adaptable entrainment across tested stimuli. These results establish a direct mechanistic link between phase sensitivity and emergent collective synchronization, demonstrating how excitation–inhibition interactions influence large-scale rhythmic coherence in firefly populations.

## Introduction

Every year as spring turns to summer, thousands of *Photuris frontalis* fireflies emerge across the southeastern United States and produce one of nature’s most striking examples of biological synchrony (*1*). Despite intrinsic variability in individual flash timing, these populations generate a coherent, population-wide bioluminescent flash pattern as they search for mates (*2, 3*). While synchronized flashing has been documented in multiple firefly species (*4–7*), the individual-level interaction rules that enable such robust synchrony remain unresolved. In particular, there are no direct empirical measurements quantifying how North American fireflies adjust their intrinsic flash timing in response to precisely timed visual perturbations (*8*). In the synchronization literature, these measurements comprise a phase response curve (PRC), which is a fundamental quantity that determines how the phase of an intrinsic oscillator shifts following external stimulation (*9, 10*). The phase response curve for several species of Southeast Asian *Pteroptyx* fireflies has been studied (*11, 12*) to various degrees of effect: Hanson and Buck found examples of excitatory-only coupling in *Pteroptyx malaccae* (*13*) while Moiseff and Copeland found that different kinds of stimulus produced different phase response effects in the same species (*12*). However, despite decades of theoretical work modeling fireflies as pulse-coupled oscillators (*8, 14, 15*), no North American firefly species has an experimentally derived PRC.

This absence represents a fundamental gap. PRCs provide a quantitative bridge between microscopic (individual-level) behavioral rules and macroscopic (population-level) synchronization dynamics, constraining entrainment, the stability of synchronous states, and the collective patterns that emerge in coupled-oscillator systems (*9, 16, 17*). Without empirical PRCs, mechanistic models necessarily assume forms of coupling strength and phase sensitivity, limiting their ability to explain how large, heterogeneous groups achieve rapid, precise synchrony in nature. This gap is especially limiting given that individual firefly flash timing is intrinsically variable within and across individuals (*18*). As a result, connecting perturbation data to mechanistic models requires high temporal resolution, repeated sampling across phases, and explicit quantification of response variability and uncertainty.

Here we address this gap by directly measuring the phase-response curve of a North American firefly species. We performed controlled perturbation assays on isolated P. frontalis males, presenting fixed-frequency light stimuli while recording flash timing with high temporal resolution. Our experiments reveal both phase *advances* and phase *delays* and enable the construction of the first data-driven PRC for any North American firefly species. The experimentally-derived PRC describes a regime of interaction dynamics most active to either side of an individual’s own action potential spike, understandable as the mechanistic influence of an impulse function upon an oscillator. We expand upon this PRC by synthesizing the phase response with empirical characterizations of endogenous oscillation and flash period variability to formulate a biologically parameterized light-controlled oscillator (LCO) model, following modeling techniques from (*19*) and (*8*). Simulations using this model reproduce key features of *P. frontalis* synchronization and demonstrate how experimentally measured phase sensitivity governs entrainment, robustness, and collective timing. Together, these results provide a mechanistic foundation for understanding synchronous flashing in *P. frontalis* and establish PRC-based modeling as a critical tool for explaining collective signal coordination in fireflies and other biological oscillators. In this paper we first outline our experimental and analytical techniques, then present results showing the period entrainment and phase response of individuals exposed to different driving frequencies. We finish with a discussion of the ramifications of this discovery.

## Experimental methods

### Field site

Field experiments took place at Congaree National Park in South Carolina. We conducted experiments each May from 2021 through 2025. To adhere to Park policies, we withhold the exact locations here but these can be provided upon direct request.

### Behavioral assays

All field experiments were performed inside a single 1.9m x 1m x 1m Alvantor Privacy Pop Up Tent covered with a 4.87m x 7.32m tarp to ensure isolation from external light sources and total darkness (Figure 1A). One GoPro Max camera recording in Hero mode at 5.6k resolution, 60 fps, and max ISO (6400) was used to capture each experiment. We applied black electrical tape on the screens and LEDs of the cameras so as not to perturb fireflies with additional artificial light signals that were not part of the experimental stimulus.

**Figure 1:**
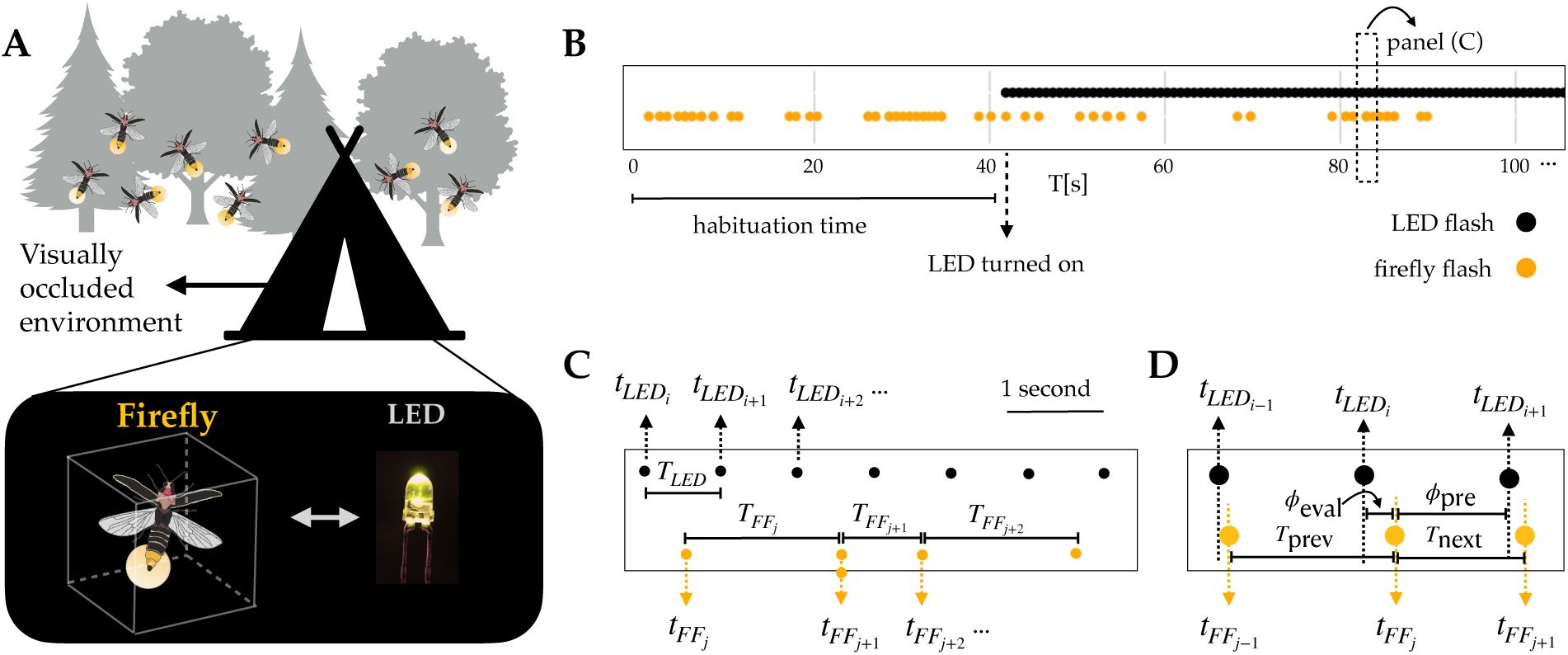
Experimental and analytical overview. (**A**) We conducted hundreds of experimental assays wherein a single firefly is enclosed in a transparent cage and exposed to an LED flashing a specific stimulus frequency. (**B**) We produce timeseries (yellow points are the firefly flashes, black points are the LED flashes) of each experiment by detecting flashes in the recordings. In each experiment, we first allow habituation to observe the firefly flashing naturally before the stimulus arrives, following techniques in (*12*). (**C**) The change in the behavior is quantified by measuring the duration between the onsets of firefly flashes (flash period) and the timing of each firefly flash relative to the timing of the stimulus LED (flash phase). (**D**) The definition of the phase variables come directly from data. *ø_eval_* is the half-wrapped *ø_pre_* and is used to contextualize phase response curve calculations to the nearest LED flash.

### Firefly behavior

Male *P. frontalis* emitted flashes continuously throughout the duration of the recordings, occasionally producing flash trains lasting several minutes, and occasionally going dark. We measured the inter-flash gap, defined as the time between the start of a flash and the end of a prior flash, for all recordings. Additionally, individual *P. frontalis* males were seen to produce flashes between 20 and 40ms in length (2-3 frames at 60fps). We measured the behavior of 127 individual fireflies in this way.

### Driven LED assay

In each experiment, one firefly was placed in a soft-sided 30cm x 30cm x 30cm RestCloud Butterfly Cage. A breadboard with one 585 nanometer wavelength LED was placed at a distance of half a meter (fifty centimeters) from the firefly. The LED was wired through 31,000 ohms of resistance in series to dim the light to the approximately same level as that of the firefly (see Supplementary Figure S1), and the flash is controlled with an Arduino Uno. Each flash of the LED was controlled with an Arduino digitalWrite method sending a signal from the micro-controller to the breadboard, dimmed by resistance. The LED was programmed to emit a square pulse 30ms in duration, then wait a specified flash period *T_LED_* before flashing again, continuously until turned off. This protocol follows precedent from (*12*) and (*20*) wherein flashing fireflies were exposed to LED stimulus after a brief self-oscillation habituation time.

Each of our experiments began with a recorded habituation time (of 70.2 ± 30.2s), letting the firefly flash with no stimulus present, to observe its natural behavior. Following this, we turned the LED on and filmed the pair for four to ten minutes (see Table S1 for more details) to observe the phase response actions of each individual firefly. At the end of each experiment, the firefly was released and a new firefly was captured and brought into the tent to initiate the next experiment.

### Analysis methods

After recording, flash positions in each video frame were extracted by applying a global pixel intensity threshold to the green channel, as in (*4*).

Because the LED position was fixed and the firefly remained confined to the butterfly cage, we draw manual bounding boxes around the two-dimensional spatial points to separate LED from firefly flashes. For each experiment, we constructed a time series of firefly and LED flashes (Figure 1B).

We addressed two main sources of noise. First, firefly attention: fireflies rarely flashed on every LED cycle, and some individuals ignored the LED entirely. To handle this, we slid a five–second window across the time series and analyzed only windows containing at least three firefly flashes, deemed to be *activity windows*. The period of the *j* th flash or the **firefly period** is

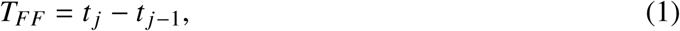

where *t _j_* and *t _j_*_–1_ are the timestamps of the starts of the *j* th and ( *j* – 1)th flashes, respectively (Figure 1C). We hereafter will use *T_FF_*to denote the firefly flash period and *T_LED_* to denote the LED flash period where appropriate. Within each valid window, the average flash period was also computed as

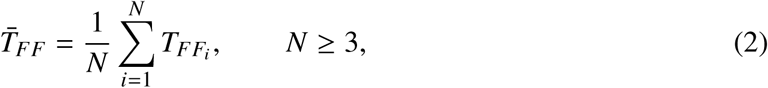

To facilitate comparison across trials, each individual’s flash train was segmented into uninterrupted **bouts**, defined as activity windows separated by less than *r_b_* = 2.0 s.

This restricts analysis to segments of time wherein the firefly behaves approximately as an oscillator. Each firefly bout is required to have at least three flashes so that at least two instances of *T_FF_* can be calculated.

The final step of constructing our dataset is to aggregate the bouts of each experiment and each LED frequency so that we can analyze phase dynamics.

### Time series analysis

For each individual and experimental population we calculate *T_FF_* and the phase of oscillation relative to the driving LED stimulus.

Given two successive firefly flashes at times *t_i_*_–1_ and *t_i_*, the **phase** of a firefly stimulus occurring at time *s* e (*t_i_*_–1_*, t_i_*) is

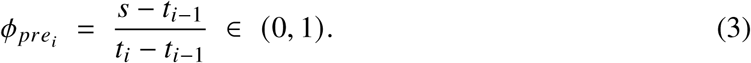

We can additionally center this in phase space around *ø* = 0 by creating the wrapped quantity *ø_eval_*:

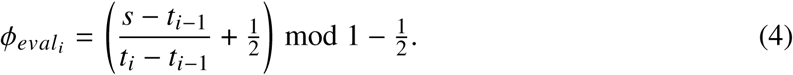

This produces a phase difference that is positive when the firefly flash occurs at a time after the nearest LED flash and is negative when the firefly flash occurs at a time before the nearest LED flash, a convenient representation of the phase for understanding the phase response dynamics (see Figure 1D for a visual description of this effect).

The phase response ratio *R*(*ø_eval_*) measures the ratio of successive periods as a way to evaluate how exposure to a stimulus changes the firefly behavior, and will be explained in detail in the following section.

### Modeling

Perturbation testing provides an experimental test bed for fitting oscillator models to firefly behavior. By applying controlled LED flashes and measuring the resulting changes in *T_FF_*, we can infer a *phase–response curve* (PRC) — the function describing how a stimulus at a given phase advances or delays the next flash.

A phase–response curve maps the phase at which a perturbation arrives to the phase shift it produces, often measured as a ratio of periods (*21*). There are several kinds of interactions that can occur, characterized by the type of the PRC. Type I PRCs advance the phase monotonically, whereas Type II PRCs have a biphasic shape with both advances and delays (*21–23*). Type II curves, common in biological and neural oscillators (*24,25*), are especially important for synchrony because they can either speed up or slow down an oscillator depending on when the pulse arrives. From our phase response curves the important quantity to measure is the impulse function *Z* (*ø*) which describes how much “kick” a stimulus provides to the oscillator in question.

We model each firefly within a bout as a phase integrator under the following definition

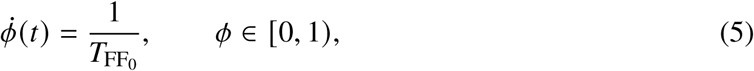

where *T*_FF0_ is the natural flash period of the firefly without stimulation. A natural flash occurs when

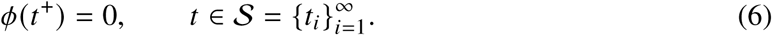

where *S* is the set of spike timings.

Because we are in the pulse-coupled paradigm, we choose to model interactions between oscillators with instantaneous phase jumps instead of continuous small influence. This leaves us in the integrate-and-fire regime, wherein a pulse (LED flash) at time *t_m_* causes an instantaneous phase jump

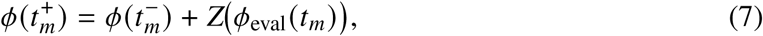

as typically defined in the literature (*26, 27*). Here *ø*(*t*^−^_m_) =: *ø*_pre_ e [0, 1) is the pre–kick phase and *ø*_eval_ is its wrapped version in the plotting domain as defined in Equation 4. If the kick crosses threshold,

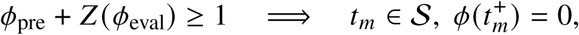

meaning an immediate flash and reset of the firefly phase *ø*_pre_ to 0. To find *Z* (*ø*_eval_) we experimentally measure

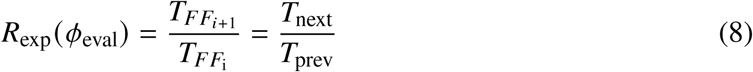

where *T*_prev_ represents the firefly period before the LED pulse and *T*_next_ represents the firefly period during the LED stimulation. This is the quantity we plot in our phase response curves in Fig 3A. If the pulse does *not* cause an immediate spike and occurs outside refractory, we can rewrite the ratio as the following:

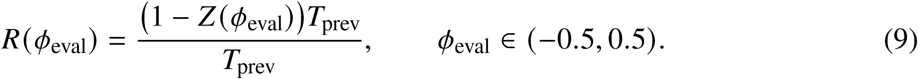

Here the numerator represents adding the impulse function *Z* to the value of the phase at the remaining fraction of the cycle. We adopt the convention *R <* 1 = phase advance (*T_next_* is shorter than *T_prev_*), which in turn implies *Z >* 0 = phase advance, from our definitions of *ø_eval_*. Therefore within the integrate–and–fire domain we recover the effective impulse PRC as

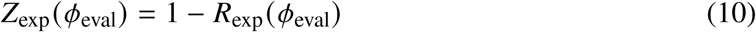

which describes the amount of influence is given to an individual perceiving a flash at the signed distance *ø*_eval_ from its own flash time.

## Results

### *Photuris frontalis* fireflies entrain to driven stimuli in predictable ways

In conditions where the LED flashes occurred shortly after the firefly flashes (early in the firefly’s next oscillatory cycle)the period was typically lengthened, consistent with inhibitory effects predicted by standard phase-resetting models (see Methods for our analysis of the phase response curve shape). Conversely, in conditions where the LED preceded the firefly flash—producing zero or positive delays—the firefly period tended to shorten, supporting excitatory interactions. Overall, the fireflies’ induced behavior appears to shift in the direction of the frequency difference between the stimulus and their intrinsic oscillation, consistent with phase-dependent entrainment dynamics. The magnitude of this entrainment effect varies depending on *T_LED_*, and in particular depending on the *difference* between the *T_LED_* and *T_FF_*_0_. We show evidence for this result with the following figures.

The accumulated distributions of *T_FF_* for treatments pre- and post-LED introduction are shown in Figure 2A.

**Figure 2:**
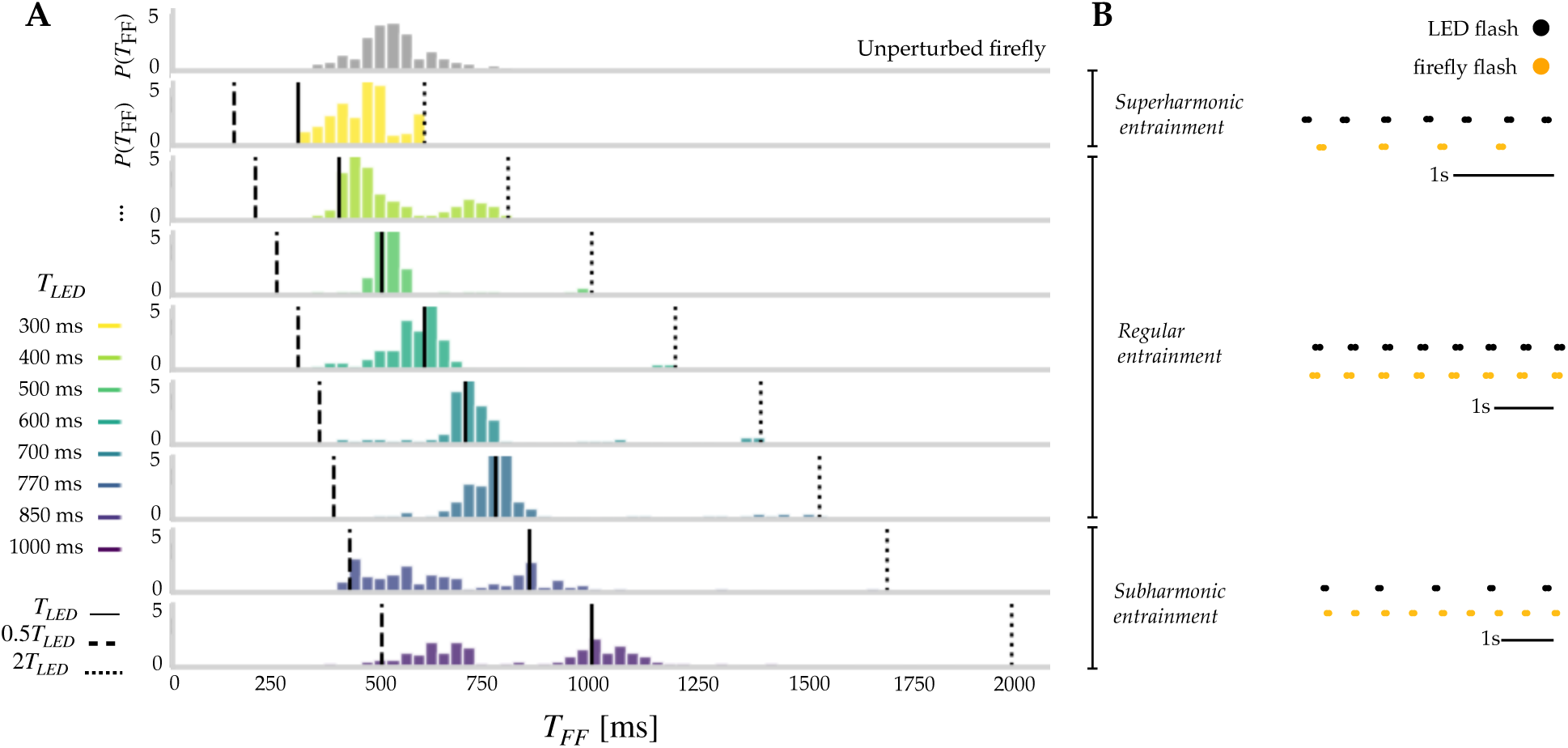
Induced flash period distributions in perturbation experiments. (**A**) Induced flash period of *Photuris frontalis* fireflies in response to each *T_LED_*. The horizontal axis is the firefly flash period *T_FF_*, measured from the end of prior flashes to the start of subsequent flashes. The vertical axis is the probability density for the occurrence of that particular *T_FF_*. The bin width for each histogram is 0.033s. **(Gray, top row)** shows the accumulated values of *T_FF_* during habituation of *N=127* individuals. **(Colored rows)** show *T_FF_* for the same individuals in the minutes of experiment wherein LED stimulus is constantly present. The three lines superimposed upon each figure represent the location of *T_LED_* (solid line), 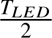 (dashed line), and 2 * *T_LED_* (dotted line). (**B**) Here we examine three different motifs of periodic response present in the data. (**Top**) Superharmonic entrainment, wherein 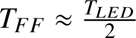, here exhibited by the 300ms driving frequency (black dots) and the synchronized response of the firefly (orange dots). In this time series, we are viewing between t=306.5s and t=309s in experiments conducted May 25th 2025. (**Middle**) Regular entrainment, wherein *T_FF_* ≈ *T_LED_*, here exhibited by an experimental timeseries with *T_LED_* = 770ms. In this time series, we are viewing between t=374s and t=380s in experiments conducted May 21st 2021. (**Bottom**) Sub-harmonic entrainment, wherein *T_FF_* ≈ 2*T_LED_*, here exhibited by an experimental timeseries with *T_LED_* = 850ms. In this time series, we are viewing between t=189s and t=193s, from experiments conducted May 28th 2021.

In four experimental conditions (*T_LED_* =300ms, 400ms, 500ms, and 600ms), the firefly’s mean period decreases after stimulation, suggesting possible phase-advance interactions from excitation (Supplementary Table S2). In the 700ms, 770ms, 850ms, and 1000ms conditions, the mean period increases, indicating phase delays, possibly from inhibition. However, looking past the mean, there are more subtle dynamics. In the 300ms condition, the median behavior was largely indistinguishable from the undisturbed behavior (Supplementary Table S2), suggesting the fireflies mostly ignored this stimulus. However, there is an increase in the mode, also represented by a small peak around twice the LED frequency (dotted lines in Figure 2 A), which suggests a marginal superharmonic entrainment effect (see Figure 2 B). These small peaks at twice the frequency also occur in the 500ms, 600ms, and 700ms conditions, albeit in a minor capacity. However, we observe a mode *decrease* in the 850ms condition, which we also see via observation of the suharmonic entrainment motif observed in Figure 2A, with a wide peak around half the LED frequency. This same subharmonic entrainment exists in the 1000ms frequency, but in both cases, we can see regular entrainment as well.

The presence of these harmonics is further confirmed when investigating the amount of flashes per cycle typically exuded by individuals when under perturbation (Supplementary Figure S2A) Additionally, across all conditions, the firefly flashed most often when the *T_LED_* closely matched the mean natural period of the firefly *T_FF_*_0_ (see Supplementary Figure S2B).

Taken together, these effects suggest that *P. frontalis* responds to the LED stimulus with both excitatory and inhibitory influences, depending on the relative timing (phase) between its intrinsic rhythm and the external stimulus. When necessary, individuals either slow down or speed up their flash period, towards whichever is closer to their normal behavior: the driving stimulus or one of its harmonics. This is confirmed by the phase response curve for each treatment, which we show in Figure 3A.

**Figure 3:**
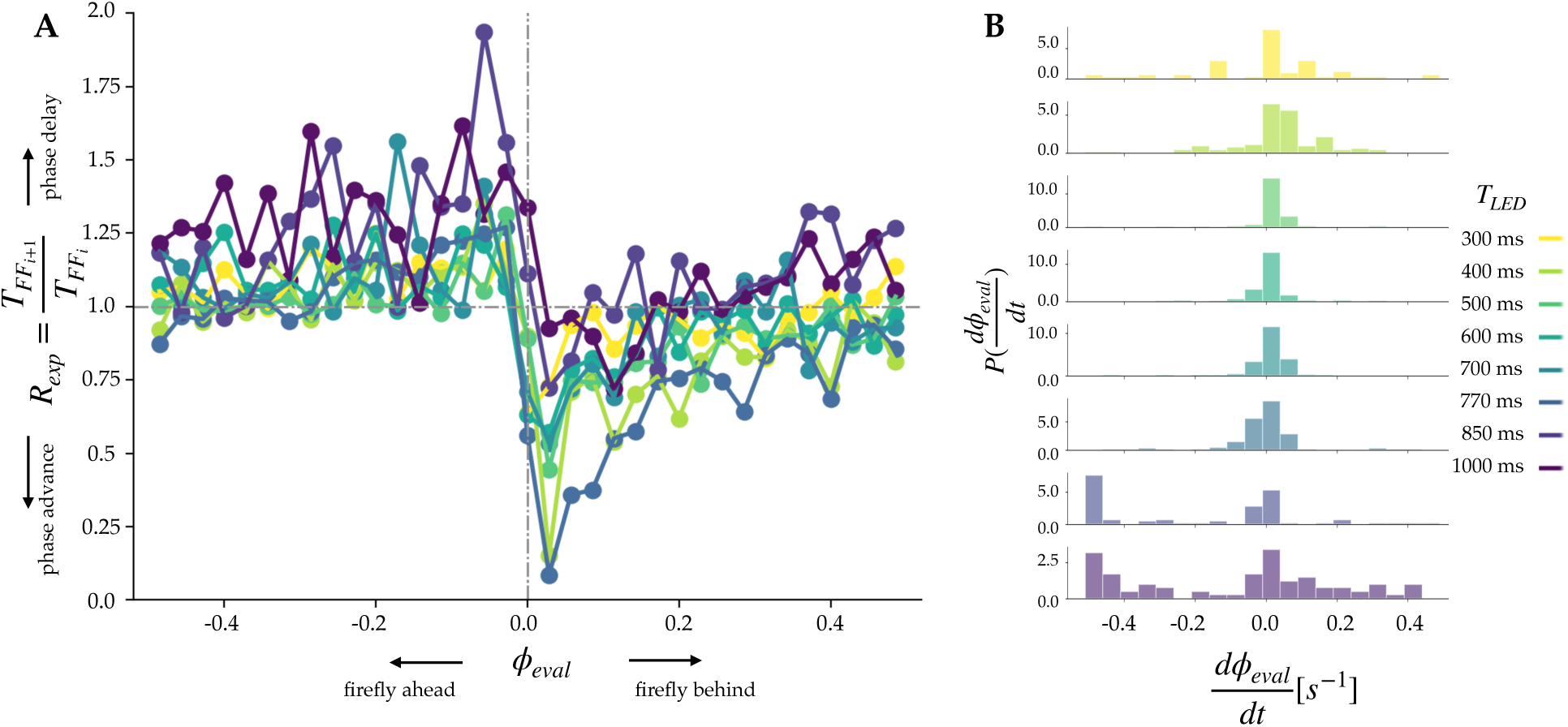
Phase response curve for *Photuris frontalis.* (**A**) Phase response curve for *Photuris frontalis* fireflies in response to a range of driving LED stimuli. Each line represents the mean flash period duration 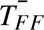 for fireflies following different driving LED stimulus frequency period (indicated by color) as a function of the LED–firefly phase difference in ms. All values taken from *bouts* of activity as defined in the methods section. Negative values of phase, *ø_eval_*, indicate the LED flash occurs early in the firefly cycle (just after a firefly flash) and positive values indicate the LED flash occurs late in the firefly cycle (just before a firefly flash). The y-axis indicates the value of R(*ø_eval_*), the ratio of the next period to the prior period of a firefly exposed to stimulus at the relative phase *phi*. (**B**) Phase derivatives relative to the firefly period are stable over bouts. For each individual in each experiment we calculate the series *ø_eval__i_*_=1_*_…n_* for each of *n* firefly flashes, then calculate the change between each. Derivatives of the time delay values near 0 indicate repeated alignment between firefly and LED flash timing, suggesting synchronization. Time delay values to the left of the dotted line indicate the firefly is flashing before the LED, on average, and vice versa for values to the right of the dotted line.

This figure demonstrates the relationship between LED-firefly phase differences and the induced *T_FF_*across multiple timing conditions, and we can see a typical type-II shape following definitions in (*24*), additionally providing evidence for these inhibitory and excitatory interactions. In particular, they seem to be the strongest when the firefly observes a flash immediately following their own flash or immediately preceding the time they were about to flash (for example, close to *ø_eval_* = 0). Furthermore, close to *ø_eval_* = 0, the amplitude of the PRC is highest when *T_LED_* is close to *T_FF_*_0_. An alternative representation of the phase response curve on domain *ø_pre_* e (0, 1] can be seen in Supplementary Figure S3B, with a notable spike (delay) in the early domain and dip (advance) in the late domain. These values correspond to the same values of *ø_eval_*represented in Figure 3A.

In addition, we observe the distribution of phase derivatives, i.e. the change in phase between the driving stimulus and the driven firefly, as a function of the LED frequency in Figure 3B. These show us that for those conditions wherein *T_FF_* matches *T_LED_*, the phase difference between the two signals also remains consistent, suggesting phase-locked, entrained, or otherwise synchronized behavior.

### Simple model simulation reproduces observed behavior

We define a simple simulation using the observed impulse function *Z* (*ø*) derived from the phase response curve as a validation technique for our results. We also note by ocular regression that the shape of the phase response curve in Figure 3A approximates that of a tanh function, and attempt to fit a parameterized tanh curve to our data. Here we describe those efforts.

### Modeling R with a parametric tanh-step function

We model the measured ratio between two consecutive firefly periods as a wrapped hyperbolic-tangent step in the evaluation phase *ø*: = *ø*_eval_ e –0.5, 0.5, then recover the corresponding impulse PRC *Z* (*ø*) using the exact integrate–and–fire (IAF) identity. Choice of this model comes from studying the literature on models that produce an appropriate shape and insights from (*28*) on nonlinear oscillators.

We posit the following parametric form for the ratio:

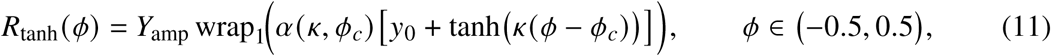

where the 1-cycle wrapping and the seam–normalization are

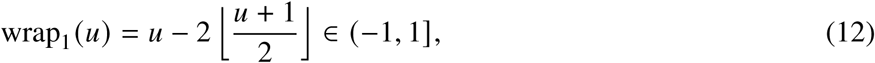

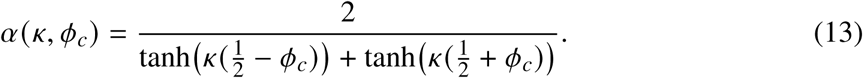

This model uses the following parameters and domains:

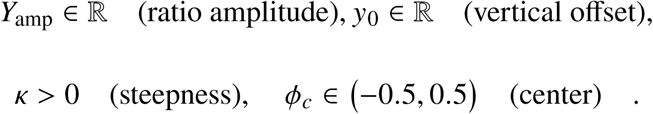

For impulses that do not cause instantaneous spiking, we can use the IAF geometry described in equations 9 and 10 to find the parameterized impulse function. Applying (11) to the fitted ratio model (10) yields the induced impulse PRC

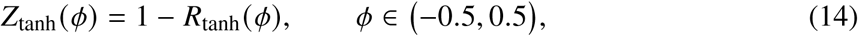

which we compare directly to the empirical *Z*_exp_(*ø*) formed from data by the same identity with *R*_exp_. Substituting and solving, we use the ratio model

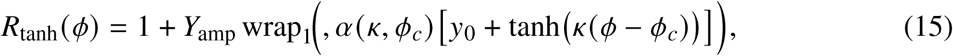

and its corresponding

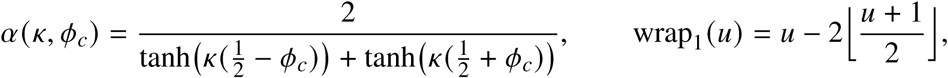

to produce the following form for Z:

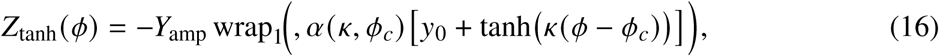

We quantify the impact of variability in the fitting parameters again by simulation. We set up a two-agent numerical model wherein one agent behaves deterministically as a LED frequency and the other behaves according to the phase response model we have selected. Each simulation runs 30,000 timesteps (reproducing five minutes of experiment time) and introduces the driving stimulus after 6000 timesteps (one minute of experiment time), reproducing the setup of our experimentation. Each simulation parameterization is simulated 100 times, and the results are aggregated and grouped by each parameter and each *T_LED_*. We evaluate the shape of the impulse function against the experimentally calculated *Z* (*ø*) in Figure 4. To test the efficacy of each model in describing the dynamics observed by *P. frontalis* interacting with a driving LED stimulus, we program simulations of the dynamics and run parameter sweeps over all parameters in Θ for each LED frequency *T_LED_*. To assess the correspondence between experimental observations and simulations of the candidate models, we evaluate the posterior distribution of firefly periods *P*(*T_FF_*) by computing the Wasserstein distance statistic between the distribution of measured *T_FF_* from simulation and experiment, for each *T_LED_*. This is an adoption of a technique widely used in goodness-of-fit testing between two distributions (*29*). The results of this can be examined in Figure 5.

**Figure 4:**
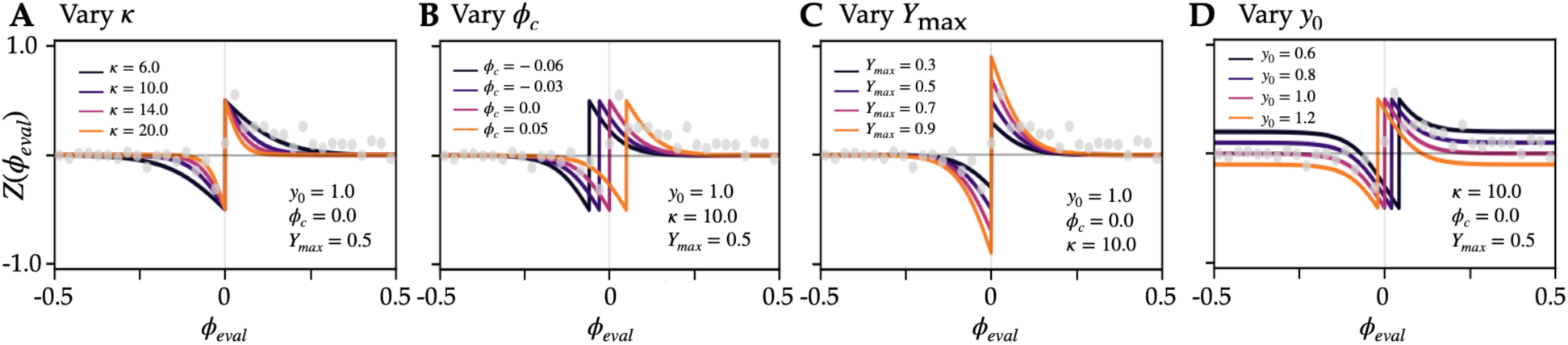
Fitting the impulse function of the PRC. Gray dots show experimental *Z* (*ø*), the impulse of the experimental PRC, calculated as 1 – *R*(*ø*). Each inset varies either the sensitivity (*н*) to light signals(**A**), the refractory period *ø_c_* (**B**), the strength (*y_max_*) of the response (**C**), and the directional bias (*y*_0_) of the first impulse (**D**). Fixed and varied parameter values are listed in the legends.

**Figure 5:**
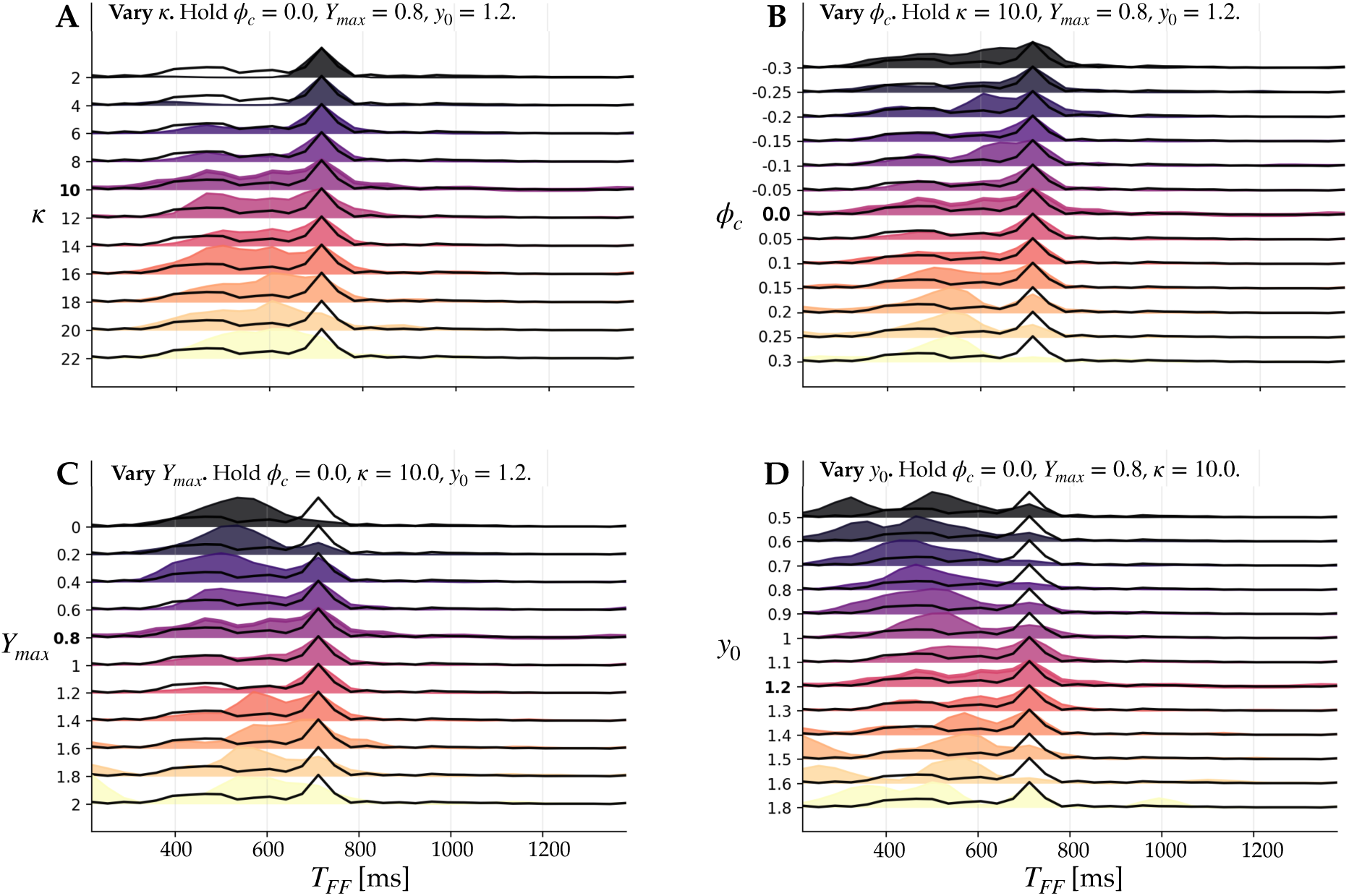
Parameter sweep variation. As an example we investigate the impact of changing the parameters of the model curve for *T_LED_* = 700ms. In all plots, the black line represents the experimental distribution of *T_FF_* for all experiments where *T_LED_* = 700ms. Each panel shows how tuning one parameter impacts the downstream distribution of *T_FF_*. Y-axis for each shows the variation tested for the parameter in question, and the value in bold is the best fit parameter over 100 simulations as measured by the Mode-Wasserstein distance (see Supplementary Methods text for distance metric definition). For this specific *T_LED_*, the best fit is *н* = 10.0*, ø_c_* = 0.0*, Y_max_* = 0.8*, y*_0_ = 1.2. For all other LED frequencies, please see Supplementary Table S3.

In Figure 5, sweeping parameters primarily reshapes the distributions of *T_FF_* by changing their spread and the relative weight of short-versus long-period components, rather than producing a uniform shift in a single characteristic timescale across all settings. Varying *н* primarily changes how sharply and strongly *Z* (*ø*_eval_) responds near the reset, making the effective phase correction more consistent across interactions and yielding tighter *T_FF_* distributions. Varying *ø_c_* mainly shifts where in phase the advance and delay parts of *Z* occur, altering when coupling is most effective within the cycle and thereby changing the relative weight of shorter versus longer periods without requiring a uniform shift of all timescales. Varying *Y*_max_ changes the magnitude of the phase kick, which broadens the statistics and increases the probability of short periods. Finally, varying *y*_0_ produces the largest qualitative changes in the shape and offset of *Z* (*ø*_eval_), modifying the balance between phase advance and delay over a wide range of phases, consistent with the broader and more variable *T_FF_* distributions observed for some *y*_0_ values in Figure 5.

In Figure 6 we compare the results of simulation to our experiments. Here we show simulations using the data directly, implementing each jump as a direct mapping from the empirical phase response curve, as well as fits from our parametric tanh function. This figure shows that a simple integrate-and-fire model with the fitted phase-resetting curve *Z* = 1 – *R* can accurately reproduce the real flashing behavior of fireflies under periodic LED stimulation.

**Figure 6:**
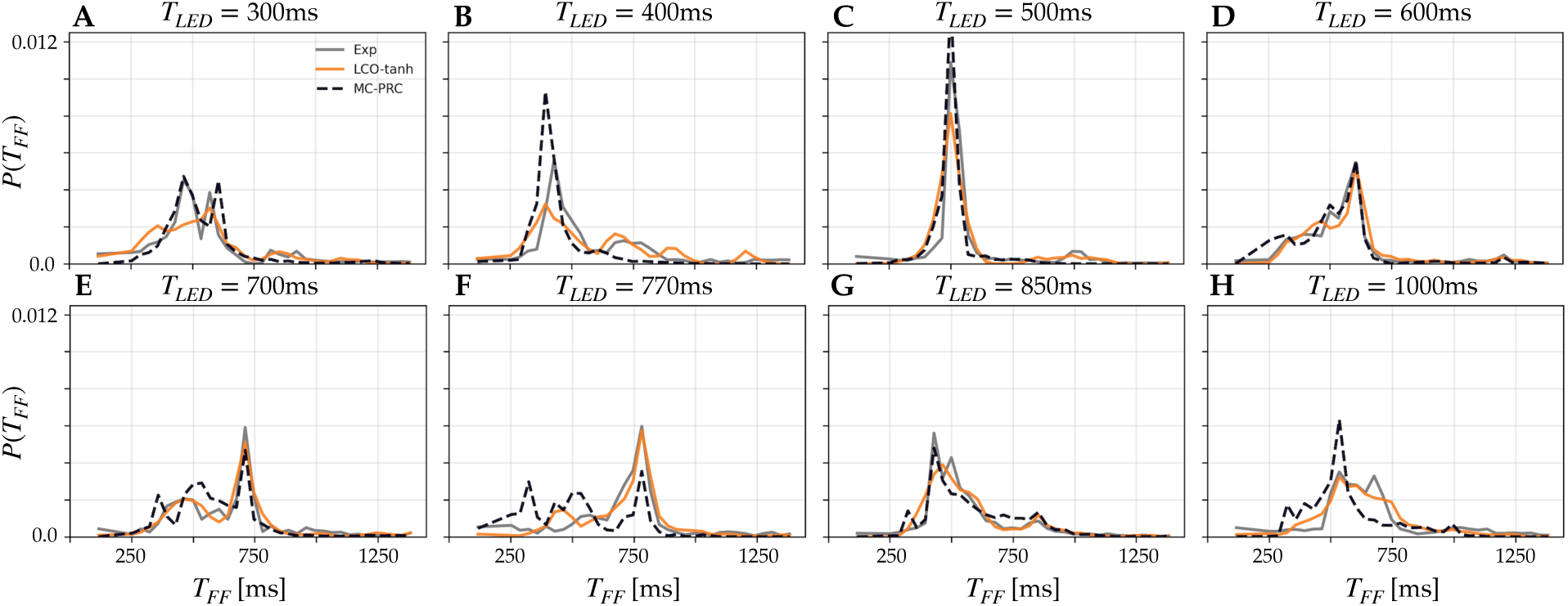
Best performing simulation results. (**A-G**) For each of the LED frequencies studied we show the experimental results (dark gray line) alongside two model simulations, from the non-parametric PRC calculated directly from experimental data (light-gray, dotted line) and the parametric tanh-shaped integrate and fire model (solid orange line), which uses *Z* (*ø*) = 1 – *R*(*ø*) as its impulse function (described in Equation 16). To match experiment, twenty trials at each frequency were simulated for 300 seconds, where an LED at the fixed frequency is exposed to a firefly oscillator. For a full description of both simulations, please see the supplementary information Materials and Methods section.

The close overlap between distribution curves not only captures the position of the main peaks but also aligns the overall shape of the probability density: this shows that the model does a good job of calculating the true timing adjustments individual fireflies make when an external pulse arrives at different phases. This means the experimentally measured phase response curve is predictive, not just descriptive. Once *Z* (*ø*) is known, we can forecast how a firefly’s intrinsic oscillator will entrain to different stimulation frequencies.

## Discussion

A measured phase response curve (PRC) collapses the complex internal dynamics of an individual firefly into an experimentally accessible causal rule: the phase-dependent transformation that maps a brief visual pulse into an advance or delay of the next flash (*10*). This reduction allows entrainment, phase-locking, and stability to be analyzed directly from behavioral data, without requiring explicit assumptions about the underlying photoneural circuitry. Our results place firefly synchronization on the same quantitative footing as canonical biological oscillators, such as neurons and circadian clocks, where PRCs serve as the bridge between individual-level physiology and population-level timing (*30*). In practical terms, the PRC specifies the expectation of entrainment under periodic forcing, the size and direction of pulse-induced phase shifts, and the parameter domains where stable locking should occur (*10*).

Decades of coupled-oscillator theory show that the sign structure and shape of the PRC constrain whether mutual excitation stabilizes in-phase synchrony, favors phase-shifted solutions, or destabilizes alignment altogether (*31*). In neural and biological oscillators, purely advancing (Type I) PRCs under strictly excitatory coupling typically disfavor robust in-phase synchrony (*10*), whereas bidirectional (Type II) PRCs, those that support excitatory and inhibitory regimes, readily support stable zero-lag locking necessary for synchronization (*10, 25, 32*). The steepness and bandwidth of the sensitive phase window further tune the size of the basin of attraction (*33*).

This empirically parsed PRC therefore does more than summarize single-animal sensitivity: it validates the existence of synchronization in this species from first principles and predicts whether visually excited individuals tend to contract or expand phase differences, and may be extended to predict whether how groups of this species might fall into clustered/chimera-like states depending on operating conditions (*2*).

Under pulse coupling, a flash from a sender updates the receiver via the phase-resetting map *ø → ø* + *Z* (*ø*), so the local slope of this map around the sampled phase window determines what happens to phase differences between the driver and oscillator. The measured PRC thus behaves as an *effective interaction kernel*, and can be scaled in any future investigation by visibility (and thus connectivity within a complex network structure) to produce all manner of emergent dynamics (*32*). In the idealized limit of identical pulse-coupled oscillators, the Mirollo–Strogatz theory ties such maps to global synchronization (*26*); moving beyond that ideal, heterogeneity in PRC amplitude or timing shifts the fixed point and can produce leader–follower asymmetries or multistability.

Parsing the PRC for *P. frontalis* supplies the missing microscopic law connecting visual pulses to timing updates, transforming qualitative descriptions of synchrony into quantitative, testable predictions for future studies. Across many biological oscillators, temperature can substantially change the intrinsic period via kinetic rate scaling, while key phase-dependent structure is comparatively preserved rather than being uniformly remapped (*34, 35*). This means that changes in ambient temperature could shift the length of phase-locked periods rather than abolishing synchrony altogether. This manifests physically as a decrease in the collective period at higher temperatures, which we observe in the field; however, it would be interesting to study this under artificially induced and controlled temperature conditions, especially those much colder than normal, to test and attempt to break the species’ entrainment capabilities. Second, embedding the measured PRC into empirically derived visibility graphs predicts a critical transition in collective order: below a critical threshold of line-of-sight connectivity, global synchrony will likely break into clustered states (*36*) or wave-like structures (*37*). A version of clustered synchronization has been observed in the field with this species (*2*) but would be interesting to measure in controlled experimental conditions to induce specific structures. Finally, species-specific asymmetries in PRC structure offer a mechanistic basis for signal segregation among sympatric firefly species, predicting resistance to heterospecific entrainment despite shared habitats. Measuring the phase response curve for predatory *Photuris lucicrescens* or *Photuris versicolor* may reveal the incapacity for biphasic entrainment: if this is the case, this flexibility may be an important evolutionary adaptation to circumvent predaceous bio-mimicry and facilitate conspecific recognition at a population level.

In the long term, implementation of PRCs into closed–loop control systems scales to real–world applications: field–deployable systems for population monitoring, conservation, and species identification recognizable by their response dynamics, bio–inspired synchronization in engineered swarms and sensor networks, become possible pursuits. Such advances could transform how we study and protect firefly biodiversity, turning behavioral signal dynamics into a quantitative and controllable tool for the physics of light-based communication, as well as ecology, evolution, and conservation technology.

## Acknowledgments

We thank the staff at Congaree National Park for facilitating field experiments and providing permits, Daniel Speiser of the University of South Carolina for measuring the light spectrum of the LED and the firefly lantern, and members of the Peleg lab for helpful discussions.

## Funding

O.P. acknowledges support from an NSF grant (2210628), and a Research Cooperation for Science Advancement grant (28219). O.P. and K.J. acknowledge support from Air Force Research Laboratory grant (FA8651-25-2-0005).

## Author contributions

O.M. contributed to project conceptualization and investigation, performed all formal analyses and methodologies, carried out all experiments, curated data, created visualizations, wrote research software, validated results, and wrote the paper draft. N.N. contributed to project investigation, wrote research software, and performed software and theoretical validation. K.J. acquired funding and lab resources, contributed to project conceptualization and investigation, and assisted with paper review and editing. O.P. supervised the project, acquired funding and lab resources, contributed to project conceptualization and investigation, and assisted with paper review and editing.

## Competing interests

There are no competing interests to declare.

## Data and materials availability

The raw timeseries of the LED and firefly behavioral assays, along with all the processing and analysis code, are available at https://github.com/peleg-lab/led_firesync.

## Supplementary materials

Materials and Methods

Supplementary Text

Figs. S1 to S3

Tables S1 to S3

## Notes

https://github.com/peleg-lab/led_firesync

### Competing Interest Statement

The authors have declared no competing interest.

## References and Notes

1. A. Moiseff, J. Copeland, A new type of synchronized flashing in a North American firefly. Journal of insect behavior 13 (4), 597–612 (2000).

2. R. Sarfati, O. Peleg, Chimera states among synchronous fireflies. Science Advances 8 (46), eadd6690 (2022), doi:10.1126/sciadv.add6690, https://www.science.org/doi/abs/10.1126/sciadv.add6690.

3. J. E. Lloyd, Bioluminescence and communication in insects. Annual review of entomology 28 (1), 131–160 (1983).

4. R. Sarfati, J. C. Hayes, Sarfati, O. Peleg, Spatiotemporal reconstruction of emergent flash synchronization in firefly swarms via stereoscopic 360-degree cameras. Royal Society Interface (2020), doi:10.1101/2020.03.19.999227.

5. R. Sarfati, L. Gaudette, J. M. Cicero, O. Peleg, Statistical analysis reveals the onset of synchrony in sparse swarms of Photinus knulli fireflies. Journal of The Royal Society Interface 19 (188), 20220007 (2022), doi:10.1098/rsif.2022.0007, https://royalsocietypublishing.org/doi/abs/10.1098/rsif.2022.0007.

6. L. Faust, A. Moiseff, J. Copeland, The night lights of Elkmont. Tenn. Conserv 64, 12–15 (1998).

7. A. Moiseff, J. Copeland, Mechanisms of synchrony in the North American firefly Photinus carolinus (Coleoptera: Lampyridae). Journal of insect behavior 8, 395–407 (1994).

8. G. M. Ramırez-Avila, J. Kurths, S. Depickere, J.-L. Deneubourg, Modeling fireflies synchronization. A mathematical modeling approach from nonlinear dynamics to complex systems pp. 131–156 (2019).

9. C. H. Johnson, Forty years of PRCs-what have we learned? Chronobiology international 16 (6), 711–743 (1999).

10. R. M. Smeal, G. B. Ermentrout, J. A. White, Phase-response curves and synchronized neural networks. Philosophical Transactions of the Royal Society B: Biological Sciences 365 (1551), 2407–2422 (2010).

11. F. E. Hanson, J. F. Case, E. Buck, J. Buck, Synchrony and flash entrainment in a New Guinea firefly. Science 174 (4005), 161–164 (1971).

12. J. Copeland, A. Moiseff, The effect of flash duration and flash shape on entrainment in Pteroptyx malaccae, a synchronic Southeast Asian firefly. Journal of insect physiology 43 (10), 965–971 (1997).

13. J. Buck, E. Buck, J. F. Case, F. E. Hanson, Control of flashing in fireflies. Journal of comparative physiology 144 (3), 287–298 (1981), doi:10.1007/BF00612560, 10.1007/BF00612560.

14. B. Ermentrout, An adaptive model for synchrony in the firefly Pteroptyx malaccae. Journal of Mathematical Biology 29 (6), 571–585 (1991).

15. D. Otte, On theories of flash synchronization in fireflies. The American Naturalist 116 (4), 587–590 (1980).

16. A. T. Winfree, Biological rhythms and the behavior of populations of coupled oscillators. Journal of theoretical biology 16 (1), 15–42 (1967).

17. Y. Kuramoto, Self-entrainment of a population of coupled non-linear oscillators, in International symposium on mathematical problems in theoretical physics (Springer) (1975), pp. 420–422.

18. O. Martin, et al., Embracing firefly flash pattern variability with data-driven species classification. Scientific Reports (2023), 10.1038/s41598-024-53671-3.

19. R. Sarfati, et al., Emergent periodicity in the collective synchronous flashing of fireflies. Elife 12, e78908 (2023).

20. A. Moiseff, J. Copeland, Firefly synchrony: A behavioral strategy to minimize visual clutter. Science 329 (2010), doi:10.1126/science.1190421.

21. A. T. Winfree, The Geometry of Biological Time (Springer, New York), 1 ed. (1980).

22. G. B. Ermentrout, D. H. Terman, Mathematical Foundations of Neuroscience, vol. 35 of Interdisciplinary Applied Mathematics (Springer, New York) (2010).

23. N. W. Schultheiss, A. A. Prinz, R. J. Butera, eds., Phase Response Curves in Neuroscience: Theory, Experiment, and Analysis (Springer, New York) (2012).

24. D. Hansel, G. Mato, C. Meunier, Synchrony in excitatory neural networks. Neural Computation 7 (2), 307–337 (1995), doi:10.1162/neco.1995.7.2.307.

25. B. Ermentrout, Type I Membranes, Phase Resetting Curves, and Synchrony. Neural Computation 8 (5), 979–1001 (1996), doi:10.1162/neco.1996.8.5.979.

26. R. E. Mirollo, S. H. Strogatz, Synchronization of pulse-coupled biological oscillators. SIAM Journal on Applied Mathematics 50 (6), 1645–1662 (1990).

27. R. Cestnik, M. Rosenblum, Inferring the phase response curve from observation of a continuously perturbed oscillator. Scientific reports 8 (1), 13606 (2018).

28. J. P. Keener, L. Glass, Global bifurcations of a periodically forced nonlinear oscillator. Journal of Mathematical Biology 21 (2), 175–190 (1984).

29. A. Ramdas, N. Garcia, M. Cuturi, Wasserstein distance and the geometry of probability distributions. Annual Review of Statistics and Its Application 4, 405–431 (2017), doi: 10.1146/annurev-statistics-060116-054114.

30. K. M. Stiefel, G. B. Ermentrout, Neurons as oscillators. Journal of neurophysiology 116 (6), 2950–2960 (2016).

31. B. Ermentrout, Y. Park, D. Wilson, Recent advances in coupled oscillator theory. Philosophical Transactions of the Royal Society A 377 (2160), 20190092 (2019).

32. A. Abouzeid, B. Ermentrout, Type-II phase resetting curve is optimal for stochastic synchrony. Physical Review E—Statistical, Nonlinear, and Soft Matter Physics 80 (1), 011911 (2009).

33. D. A. Wiley, S. H. Strogatz, M. Girvan, The size of the sync basin. Chaos: An Interdisciplinary Journal of Nonlinear Science 16 (1) (2006).

34. W. Soofi, et al., Phase maintenance in a rhythmic motor pattern during temperature changes in vivo. Journal of Neurophysiology 111 (12), 2603–2613 (2014).

35. P. François, N. Despierre, E. D. Siggia, Adaptive temperature compensation in circadian oscillations. PLoS Computational Biology 8 (7), e1002585 (2012).

36. P. S. Skardal, E. Ott, J. G. Restrepo, Cluster synchrony in systems of coupled phase oscillators with higher-order coupling. Physical Review E—Statistical, Nonlinear, and Soft Matter Physics 84 (3), 036208 (2011).

37. K. L. Rodrigues, R. Dickman, Synchronization of discrete oscillators on ring lattices and small-world networks. Journal of Statistical Mechanics: Theory and Experiment 2020 (4), 043406 (2020).

38. V. M. Panaretos, Y. Zemel, Statistical aspects of Wasserstein distances. Annual review of statistics and its application 6 (1), 405–431 (2019).

