## Supplementary Information for "Excitation–inhibition interactions mediate firefly flash synchronization"

### Supplementary Materials for
### Excitation–inhibition interactions mediate firefly flash synchronization

Owen L. Martin[1*], Nataliya Nechyporenko[1], Kaushik Jayaram[1,2]Orit Peleg[1,3,4]

[1]Department of Computer Science, University of Colorado, Boulder, CO, USA. [2]Department of Bioengineering, Imperial College, London, UK. [3]Santa Fe Institute, NM, USA. *Corresponding author.

**This PDF file includes:**

Materials and Methods

Supplementary Text

Figures S1 to S3

Tables S1 to S3

#### Materials and Methods

##### Simulation of the Light-Controlled Oscillator (LCO) Model

To characterize how different mechanistic phase–response curve (PRC) shapes generate the experimentally observed flash period distributions under periodic LED forcing, we simulated a nonlinear light-controlled oscillator (LCO) for a four-dimensional grid of model parameters. The model tracks a continuous phase variable $\phi(t) \in [0, 1)$ which increases at a rate determined by the intrinsic ("free-running") period $T_0$. For each trial, $T_0$ was sampled from the empirical distribution of pre-stimulus periods measured from isolated individuals.

Between LED stimuli, the oscillator evolves according to

$$\frac{d\phi}{dt} = \frac{1}{T_0}, \qquad \phi(t) = 1 \;\Rightarrow\; \phi \leftarrow \phi - 1.$$

LED pulses were delivered at fixed periods $p$ (in ms), as in the behavioral assays, with randomized onset time $t_{\text{start}} \in [58, 60]$ s to prevent alignment artifacts. All LED times were precomputed for a 300 s simulation window.

At each LED arrival time $t_{\text{LED}}$, the phase undergoes an instantaneous shift

$$\phi(t^+) = \phi(t^-) + \Delta\phi(\phi^-; \kappa, \phi_c, Y_{\max}, y_0),$$

where $(\kappa, \phi_c, Y_{\max}, y_0)$ parameterize a smooth sigmoidal impulse function. If $\phi(t^+) \geq 1$, the LED immediately triggers a spike.

For each LED period $p$, we evaluated all combinations of

$$\kappa \in K, \qquad \phi_c \in C, \qquad Y_{\max} \in M, \qquad y_0 \in Y_0,$$

resulting in $3 \times 5 \times 3 \times 3 = 135$ parameter sets per LED. Each parameter set was simulated for 100 independent trials. Flash periods after LED onset were binned using the same bin edges (200–1400 ms) as the experimental data.

Model fit was quantified using the Wasserstein-1 distance:

$$D_{\text{W}} = W_1(h_{\text{sim}}, h_{\text{exp}}).$$

The globally optimal parameter set was selected by minimizing $D_{\text{W}}$, with 2D parameter landscape slices computed for visualization.

#### Simulation of the Empirical PRC Model (Monte Carlo Approach)

To complement the parametric LCO model, we simulated oscillators driven directly by the experimentally measured PRCs. These empirical PRCs provide a nonparametric benchmark capturing the direct stimulus–response relationship observed in the behavioral assays.

For each LED period $p$, we computed the phase $\phi$ at which each LED pulse occurred within the firefly's free-running oscillatory cycle, defined as the fractional position between the preceding and subsequent spontaneous flashes. Each pulse therefore produced a phase–response pair $(\phi,\ T_{\text{next}}/T_{\text{prev}})$ (see Eq. 8 definitions for $T_{\text{next}}$ and $T_{\text{prev}}$). These phase–response measurements were grouped into uniform phase bins $\phi_i$, and the corresponding bin-averaged ratios,

$$R(\phi) = \frac{T_{\text{FF}_{i+1}}}{T_{\text{FF}_i}} = \frac{T_{\text{next}}}{T_{\text{prev}}},$$

yielded the empirical PRC lookup table $\{(\phi_i, r_i)\}$ used in the nonparametric simulation framework.

The firefly oscillator evolves with $\dot{\phi} = 1/T_0$ and undergoes the empirical PRC phase update following an LED pulse:

$$\phi(t^+) = \phi(t^-) + \Delta(\phi), \qquad \Delta(\phi) = 1 - R(\phi),$$

consistent with the experimental definition of $R(\phi)$.

To incorporate uncertainty in the empirical phase response curve into the nonparametric simulation, we performed Monte Carlo resampling of the phase–response measurements. Each empirical PRC consists of phase bins $\phi_i$ paired with mean $T_{FF}$ ratios $r_i$ and associated standard errors of the mean (SEM) calculated from the observed data. In each Monte Carlo realization, we constructed a perturbed PRC by replacing every $r_i$ with

$$r_i^{(k)} \sim \text{Uniform}(r_i - \text{SEM}_i,\ r_i + \text{SEM}_i),$$

thereby generating a plausible PRC consistent with the experimental uncertainty in that bin. This procedure preserves the measured phase structure of the PRC while allowing for stochastic variation in ratio magnitude.

For each $T_{LED}$, we generated 50 such perturbed PRC realizations, and for each realization we ran 100 independent oscillator simulations. The resulting ensemble of simulated flash period distributions reflects both intrinsic biological variability and uncertainty in the empirical PRC estimation, and the best Monte Carlo instance can be calculated following the statistical test framework described below.

##### Statistical Comparison Between Models and Experiment

To quantify how well each simulated oscillator reproduced the experimentally observed flash period distributions, we employed a peak-centered distance metric that captures both the accuracy of the locking period and the local agreement in distributional shape around the dominant mode. Let $h_{\text{sim}}$ and $h_{\text{exp}}$ denote the simulated and experimental flash period histograms evaluated over a shared set of bin centers $\{c_j\}$, and let $m_{\text{sim}}$ and $m_{\text{exp}}$ denote their respective modes (the bin centers at which each histogram attains its maximum).

**Mode difference.** The deviation in locking period was quantified as the absolute difference between the simulated and experimental modes,

$$\Delta m = \left|m_{\text{sim}} - m_{\text{exp}}\right| \quad \text{(ms)}.$$

To assess the similarity of histogram shapes near the locking regime, we extracted a symmetric window of width $W$ (set to $W = 200$ ms) centered on the experimental mode. Let

$$\mathcal{J} = \{j : |c_j - m_{\text{exp}}| \leq W/2\}$$

index the bins within this window. The experimental and simulated histograms were restricted to this window and renormalized to form local probability distributions,

$$p_j = \frac{h_{\text{exp}}(c_j)}{\sum_{k\in\mathcal{J}} h_{\text{exp}}(c_k)}, \qquad q_j = \frac{h_{\text{sim}}(c_j)}{\sum_{k\in\mathcal{J}} h_{\text{sim}}(c_k)}, \qquad j \in \mathcal{J}.$$

The local Wasserstein–1 distance was computed using the discrete formula (*38*),

$$W_1^{\text{local}} = \sum_{j\in\mathcal{J}} \left|F_p(c_j) - F_q(c_j)\right| \Delta c,$$

where $F_p$ and $F_q$ are the empirical cumulative distributions of $p_j$ and $q_j$, and $\Delta c$ is the bin width in milliseconds. This yields a shape-sensitive measure of local distributional disagreement, expressed in the same physical units as the mode difference.

Overall agreement between simulation and experiment was quantified by

$$D_{\text{peak}} = \sqrt{\Delta m^2 + \left(\lambda\, W_1^{\text{local}}\right)^2},$$

where $\lambda$ is a dimensionless weighting factor ($\lambda = 0.5$ in all analyses). This metric preserves the interpretability of the peak difference $\Delta m$ while incorporating a localized comparison of distributional shape near the locking peak. Best-fit parameters for both the LCO model and the empirical-PRC Monte Carlo simulations were obtained by minimizing $D_{\text{peak}}$ for each LED driving period.

For each LED period $p$, we report (i) the optimal LCO parameters and corresponding simulated flash period distributions, (ii) the best empirical-PRC Monte Carlo result, and (iii) overlay plots comparing experimental, LCO-simulated, and empirical-PRC-simulated IFI histograms.

#### Supplementary Text

##### LED Light spectra

We compared the intensities and wavelengths of the spectra of an LED pulse to a pulse of light emitted by a male *Photuris frontalis* in Figure S3. We measured the absolute irradiance (integrated from 400 to 700 nm) of firefly flashes and the LED using a spectrometer system with components from Ocean Optics (Dunedin, FL), including a SR6VN500-100 spectrometer, a QP400-1-UV-VIS optical fiber, and a CC-3 cosine corrector. We calibrated the absolute spectral response of the spectrometer using an HL-3P-CAL Vis-NIR calibrated light source and we operated the spectrometer using OceanView 2.0.20 software. When recording firefly flashes and the LED, we used integration times of 25 ms and 500 ms, respectively. At a recording distance of 1 cm, the spectral peaks and irradiance values of firefly flashes and the LED were similar: the LED had a peak spectral output of between 584 and 585 nanometers, with an irradiance of $2.22 \times 10^{13}$ photons/cm$^2$/sec, while the firefly flashes had a peak spectral output between 570 and 571 nanometers with variable irradiance values at the same order of magnitude (with the irradiance value of the flash shown in Fig. S3 being $4.23 \times 10^{13}$ photons/cm$^2$/sec). We thank our collaborator, Daniel Speiser at the University of South Carolina, for these measurements.

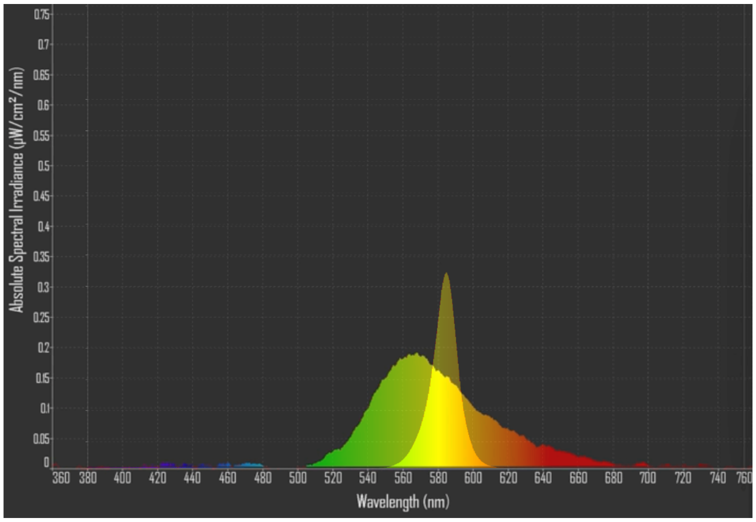


**Figure S1**: **Light spectra comparison between LED stimulus and firefly signals.** Comparison between the intensity and wavelength of the firefly spectra and the spectra of our resistance-dimmed LED system.

##### Activity variability

One aspect of the behavioral variability we wanted to capture between assay treatments is the overall activity (flashes) induced by the LED stimulus. This looks like counting the total amount of flashes in the five minute time window following the introduction of the stimulus. We can additionally *normalize* this quantity by the period of the LED pulse, under the hypothesis that all other things kept equal, more flashes seen induces more flash activity. We share these two lenses into the data in Figure S2.

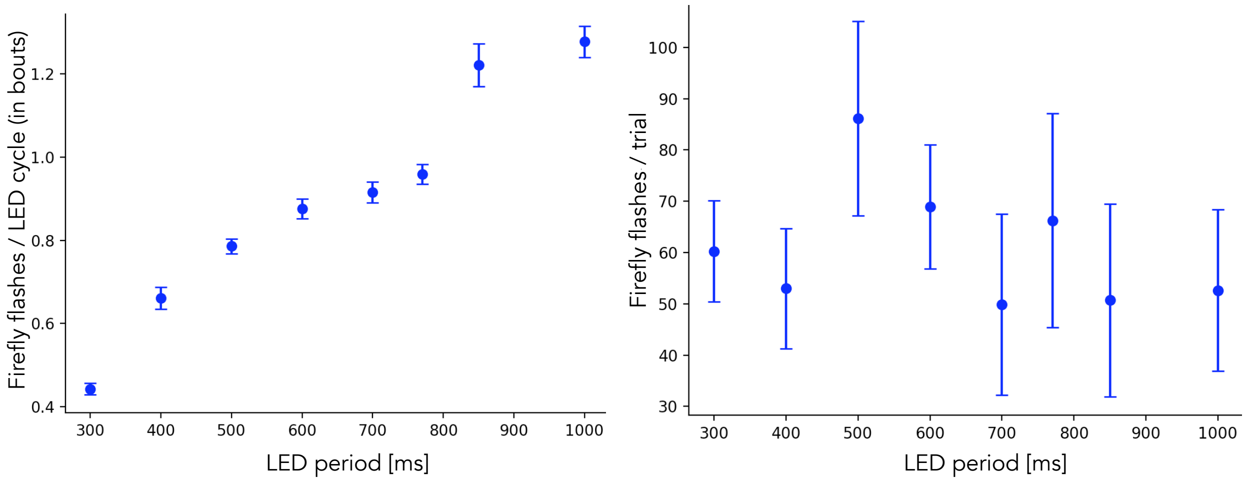


**Figure S2**: **Total activity levels per experimental condition.** Comparison between the per-cycle flash counts (left) and total flash counts (right) per each LED treatment. We see further evidence for the different entrainment regimes in the left hand plot, where each trial's induced flash count is divided by the cycle count to determine how many flashes are occuring on average during each $T_{LED}$. Below 500ms the induced flash activity is low, suggesting flashes on every other or every third beat. Between 500ms and 770ms the induced flash activity is near one flash per beat. And at 850ms and above the induced flash activity surpasses one flash per beat. The right hand plot confirms that the 500ms experimental condition, the closest to the mean behavior of *Photuris frontalis* in Congaree NP, induces the most overall activity.

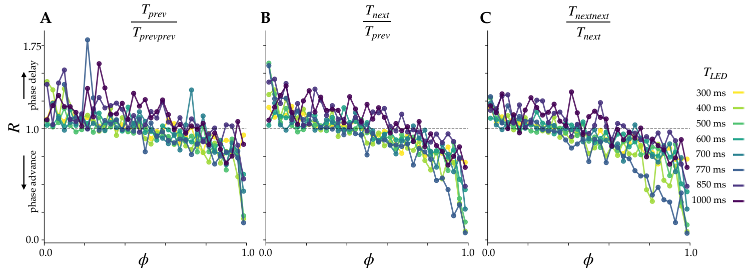


**Figure S3**: **Phase response curves on the 0-1 domain.** In (**A**) and (**C**) we examine backward-only and forward-only evaluations of the phase response curve, taking advantage of the periodicity of the driving signal, to see how stable and consistent the phase response reactions are. In (**B**) we show simply the same phase response curve as Figure 3A but unwrapped onto the 0-1 phase domain.

**Table S1**: Experimental Metadata. All $T_{\text{LED}}$ values in milliseconds (ms). $t_{\text{pre}}$ and $t_{\text{post}}$ represent experimental durations, in seconds, before (pre) the LED is introduced and while the LED and firefly (post) are flashing together.

| yyyy | mm | dd | $T_{\text{LED}}$ | °C | $t_{\text{pre}}$ | $t_{\text{post}}$ | yyyy | mm | dd | $T_{\text{LED}}$ | °C | $t_{\text{pre}}$ | $t_{\text{post}}$ |
|---|---|---|---|---|---|---|---|---|---|---|---|---|---|
| 2022 | 05 | 29 | 300 | 22.9 | 100.0 | 239.7 | 2022 | 05 | 29 | 300 | 23.1 | 40.0 | 165.6 |
| 2022 | 05 | 29 | 300 | 23.2 | 8.13 | 98.7 | 2022 | 05 | 29 | 300 | 23.6 | 63.57 | 355.2 |
| 2022 | 05 | 29 | 300 | 23.8 | 105.0 | 294.9 | 2022 | 05 | 29 | 300 | 23.9 | 60.0 | 159.9 |
| 2022 | 05 | 29 | 300 | 24.0 | 57.57 | 259.8 | 2023 | 05 | 23 | 300 | 20.1 | 145.67 | 263.1 |
| 2023 | 05 | 23 | 300 | 20.3 | 32.97 | 205.8 | 2023 | 05 | 23 | 300 | 20.4 | 123.9 | 300.6 |
| 2025 | 05 | 20 | 300 | 23.0 | 35.05 | 549.65 | 2025 | 05 | 20 | 300 | 23.4 | 56.91 | 413.41 |
| 2025 | 05 | 20 | 300 | 23.5 | 45.76 | 552.45 | 2025 | 05 | 20 | 300 | 24.0 | 45.16 | 588.76 |
| 2025 | 05 | 20 | 300 | 24.1 | 47.32 | 516.45 | 2025 | 05 | 20 | 300 | 24.2 | 54.69 | 480.15 |
| 2025 | 05 | 20 | 300 | 24.3 | 63.8 | 464.93 | 2025 | 05 | 20 | 300 | 24.4 | 52.4 | 494.11 |
| 2025 | 05 | 20 | 300 | 24.4 | 55.81 | 442.28 | 2022 | 05 | 29 | 400 | 22.9 | 57.57 | 137.2 |
| 2022 | 05 | 29 | 400 | 23.2 | 100.0 | 314.0 | 2022 | 05 | 29 | 400 | 23.2 | 60.0 | 323.2 |
| 2022 | 05 | 29 | 400 | 23.3 | 86.67 | 279.6 | 2022 | 05 | 29 | 400 | 23.4 | 60.0 | 329.2 |
| 2022 | 05 | 29 | 400 | 23.5 | 89.5 | 238.5 | 2023 | 05 | 23 | 400 | 20.1 | 44.9 | 625.6 |
| 2023 | 05 | 23 | 400 | 20.3 | 122.8 | 311.2 | 2023 | 05 | 23 | 400 | 20.4 | 60.0 | 544.0 |
| 2025 | 05 | 19 | 400 | 22.4 | 54.04 | 666.63 | 2025 | 05 | 19 | 400 | 22.5 | 55.54 | 486.03 |
| 2025 | 05 | 19 | 400 | 22.6 | 48.47 | 556.2 | 2025 | 05 | 19 | 400 | 25.6 | 61.08 | 527.87 |
| 2021 | 05 | 25 | 500 | 23.4 | 67.9 | 597.32 | 2021 | 05 | 25 | 500 | 23.7 | 125.23 | 541.29 |
| 2021 | 05 | 27 | 500 | 23.2 | 57.4 | 596.82 | 2021 | 05 | 27 | 500 | 24.1 | 70.47 | 595.81 |
| 2021 | 05 | 27 | 500 | 24.6 | 75.27 | 590.81 | 2021 | 05 | 27 | 500 | 24.9 | 65.02 | 599.82 |
| 2021 | 05 | 27 | 500 | 24.1 | 140.74 | 525.78 | 2021 | 05 | 27 | 500 | 23.4 | 66.18 | 599.82 |
| 2021 | 05 | 27 | 500 | 24.6 | 68.1 | 596.82 | 2021 | 05 | 27 | 500 | 24.9 | 69.52 | 592.81 |
| 2022 | 05 | 19 | 500 | 24.5 | 93.33 | 292.0 | 2022 | 05 | 21 | 500 | 22.1 | 50.13 | 243.5 |
| 2022 | 05 | 24 | 500 | 25.8 | 72.17 | 256.0 | 2022 | 05 | 24 | 500 | 26.1 | 31.97 | 207.5 |
| 2022 | 05 | 26 | 500 | 23.1 | 166.67 | 305.52 | 2022 | 05 | 26 | 500 | 23.2 | 60.68 | 382.52 |
| 2022 | 05 | 26 | 500 | 23.3 | 84.25 | 193.02 | 2022 | 05 | 26 | 500 | 23.4 | 16.67 | 483.02 |
| 2021 | 05 | 25 | 600 | 22.7 | 64.92 | 598.52 | 2021 | 05 | 25 | 600 | 23.0 | 66.25 | 598.52 |
| 2021 | 05 | 25 | 600 | 24.4 | 67.83 | 598.52 | 2021 | 05 | 25 | 600 | 24.4 | 63.33 | 603.32 |
| 2021 | 05 | 25 | 600 | 23.9 | 67.32 | 598.52 | 2021 | 05 | 26 | 600 | 23.9 | 66.93 | 599.72 |
| 2021 | 05 | 26 | 600 | 24.2 | 68.13 | 597.92 | 2021 | 05 | 26 | 600 | 25.6 | 69.15 | 597.32 |
| 2021 | 05 | 26 | 600 | 23.4 | 152.69 | 513.27 | 2021 | 05 | 26 | 600 | 24.4 | 69.43 | 596.72 |
| 2022 | 05 | 18 | 600 | 22.2 | 74.0 | 328.8 | 2022 | 05 | 18 | 600 | 22.3 | 72.93 | 237.6 |
| 2022 | 05 | 18 | 600 | 22.3 | 87.67 | 327.6 | 2022 | 05 | 19 | 600 | 24.2 | 22.03 | 192.6 |
| 2022 | 05 | 20 | 600 | 24.4 | 93.17 | 216.4 | 2022 | 05 | 25 | 600 | 25.6 | 64.77 | 418.82 |
| 2022 | 05 | 26 | 600 | 23.4 | 142.05 | 356.42 | 2021 | 05 | 22 | 700 | 20.4 | 69.0 | 590.81 |
| 2021 | 05 | 22 | 700 | 20.8 | 69.92 | 596.72 | 2021 | 05 | 22 | 700 | 21.4 | 64.6 | 599.22 |
| 2021 | 05 | 22 | 700 | 21.4 | 59.73 | 594.61 | 2021 | 05 | 22 | 700 | 23.2 | 63.08 | 597.42 |
| 2021 | 05 | 23 | 700 | 23.2 | 68.05 | 597.82 | 2021 | 05 | 23 | 700 | 23.7 | 56.68 | 607.62 |
| 2021 | 05 | 23 | 700 | 23.0 | 68.67 | 595.01 | 2021 | 05 | 23 | 700 | 23.8 | 95.85 | 567.02 |
| 2021 | 05 | 24 | 700 | 25.4 | 64.1 | 602.32 | 2021 | 05 | 25 | 700 | 23.8 | 10.89 | 647.14 |
| 2021 | 05 | 25 | 700 | 20.4 | 78.52 | 585.51 | 2022 | 05 | 18 | 700 | 22.2 | 41.2 | 275.1 |
| 2022 | 05 | 18 | 700 | 22.3 | 16.93 | 67.9 | 2022 | 05 | 19 | 700 | 24.7 | 60.0 | 306.6 |
| 2022 | 05 | 19 | 700 | 24.8 | 40.0 | 239.4 | 2022 | 05 | 20 | 700 | 24.6 | 53.7 | 238.7 |
| 2022 | 05 | 25 | 700 | 22.2 | 46.6 | 312.92 | 2021 | 05 | 21 | 770 | 19.9 | 30.63 | 571.35 |
| 2021 | 05 | 21 | 770 | 20.4 | 115.52 | 544.71 | 2021 | 05 | 21 | 770 | 20.8 | 87.19 | 486.33 |
| 2021 | 05 | 21 | 770 | 20.2 | 60.17 | 665.82 | 2021 | 05 | 21 | 770 | 20.8 | 40.74 | 624.4 |
| 2021 | 05 | 21 | 770 | 24.2 | 68.85 | 579.97 | 2021 | 05 | 24 | 770 | 24.1 | 55.68 | 604.27 |
| 2021 | 05 | 24 | 770 | 24.7 | 71.44 | 592.95 | 2021 | 05 | 28 | 770 | 24.7 | 66.17 | 598.78 |
| 2021 | 05 | 28 | 770 | 22.2 | 84.79 | 581.44 | 2022 | 05 | 18 | 770 | 22.2 | 103.83 | 312.8 |
| 2022 | 05 | 18 | 770 | 22.2 | 78.7 | 227.7 | 2022 | 05 | 18 | 770 | 23.1 | 30.0 | 417.07 |
| 2021 | 05 | 23 | 850 | 23.3 | 87.91 | 578.31 | 2021 | 05 | 23 | 850 | 23.4 | 107.32 | 558.75 |
| 2021 | 05 | 23 | 850 | 24.0 | 78.02 | 587.66 | 2021 | 05 | 23 | 850 | 24.0 | 97.05 | 568.1 |
| 2021 | 05 | 24 | 850 | 24.5 | 64.47 | 599.57 | 2021 | 05 | 24 | 850 | 24.0 | 178.66 | 487.31 |
| 2021 | 05 | 27 | 850 | 23.4 | 79.24 | 582.56 | 2021 | 05 | 28 | 850 | 24.3 | 68.17 | 597.87 |
| 2021 | 05 | 28 | 850 | 24.6 | 77.16 | 589.36 | 2021 | 05 | 28 | 850 | 24.6 | 73.82 | 590.21 |
| 2025 | 05 | 18 | 850 | 23.4 | 55.82 | 419.75 | 2025 | 05 | 18 | 850 | 23.5 | 57.06 | 412.03 |
| 2025 | 05 | 18 | 850 | 23.6 | 36.42 | 432.67 | 2025 | 05 | 18 | 850 | 23.7 | 58.78 | 435.25 |
| 2025 | 05 | 18 | 850 | 23.7 | 58.83 | 410.29 | 2021 | 05 | 29 | 1000 | 23.8 | 68.18 | 597.32 |
| 2021 | 05 | 29 | 1000 | 24.1 | 71.29 | 593.31 | 2021 | 05 | 29 | 1000 | 24.9 | 59.38 | 599.32 |
| 2021 | 05 | 29 | 1000 | 24.3 | 62.26 | 598.32 | 2021 | 05 | 29 | 1000 | 23.8 | 67.93 | 598.32 |
| 2021 | 05 | 29 | 1000 | 24.1 | 68.6 | 597.32 | 2021 | 05 | 29 | 1000 | 24.9 | 82.09 | 584.31 |
| 2022 | 05 | 26 | 1000 | 23.3 | 37.62 | 329.02 | 2025 | 05 | 22 | 1000 | 22.6 | 53.1 | 546.5 |
| 2025 | 05 | 22 | 1000 | 22.6 | 53.72 | 586.9 | 2025 | 05 | 22 | 1000 | 22.7 | 46.2 | 586.91 |
| 2025 | 05 | 22 | 1000 | 22.7 | 54.67 | 526.29 | 2025 | 05 | 22 | 1000 | 22.8 | 10.44 | 618.22 |
| 2025 | 05 | 22 | 1000 | 22.8 | 50.14 | 584.9 | | | | | | | |

**N**: 300 (**19**), 400 (**13**), 500 (**18**), 600 (**17**), 700 (**18**), 770 (**13**), 850 (**15**), 1000 (**14**), All (**127**)

**Table S2**: **Aggregate pre- and post-perturbation statistics for experiments. All units ms.**

| $T_{\mathrm{LED}}$ | Mean $T_{FF}$ | | Median $T_{FF}$ | | Mode $T_{FF}$ | |
|---|---|---|---|---|---|---|
| | Pre | Post | Pre | Post | Pre | Post |
| 300 | 577 ± 258 | 455 ± 74.7 | 500 | 466 | 434 | 600 |
| 400 | 642 ± 235 | 517 ± 123 | 567 | 467 | 567 | 433 |
| 500 | 548 ± 209 | 542 ± 104 | 533 | 517 | 533 | 500 |
| 600 | 616 ± 251 | 607 ± 148 | 567 | 600 | 567 | 600 |
| 700 | 615 ± 197 | 735 ± 167 | 567 | 717 | 517 | 717 |
| 770 | 607 ± 244 | 815 ± 210 | 534 | 767 | 517 | 767 |
| 850 | 510 ± 175 | 735 ± 282 | 484 | 667 | 467 | 434 |
| 1000 | 592 ± 240 | 898 ± 320 | 550 | 925 | 517 | 1033 |

**Table S3**: Best-fit phase response curve parameters for each LED period. Here, $Y_{\max}$ is the maximum phase advance, $Y_0$ the baseline offset, $\phi_c$ the phase of maximal sensitivity, and $\kappa$ controls the sharpness of the transition.

| $T_{\mathrm{LED}}$ (ms) | $Y_{max}$ | $Y_0$ | $\phi_c$ | $\kappa$ |
|---|---|---|---|---|
| 300 | 1.40 | 1.00 | +0.00 | 8.0 |
| 400 | 0.60 | 1.20 | -0.30 | 12.0 |
| 500 | 1.20 | 1.80 | -0.30 | 10.0 |
| 600 | 1.20 | 1.80 | -0.30 | 8.0 |
| 700 | 0.80 | 1.20 | +0.00 | 10.0 |
| 770 | 1.00 | 1.40 | +0.30 | 4.0 |
| 850 | 1.00 | 1.00 | +0.00 | 12.0 |
| 1000 | 1.20 | 1.20 | +0.15 | 10.0 |
